# Computational and experimental studies of the breathing motion of a protein loop: implications in *Pf*AMA1-*Pf*RON2 late-stage binding event

**DOI:** 10.1101/2021.10.10.463826

**Authors:** Suman Sinha, Anamika Biswas, Jagannath Mondal, Kalyaneswar Mandal

**Author notes:** **Corresponding Author** (J. Mondal), (K. Mandal). These authors contributed equally (Suman Sinha, Anamika Biswas).

## Abstract

Protein-protein interactions are important targets for various drug discovery campaigns. One such promising and therapeutically pertinent protein-protein complex is *Pf*AMA1-*Pf*RON2 involved in malarial parasite invasion into human red blood cells. A thorough understanding of the interactions between these macromolecular binding partners is crucial for designing better therapeutics against this age-old disease. Although crystal structures of several *Pf*AMA1-*Pf*RON2 complexes are available, the mechanism of how domain II loop associates with *Pf*RON2 is not clear. The current work investigates how the domain II loop of *Pf*AMA1 exerts its effect on the alpha helix of the *Pf*RON2, thus influencing the overall kinetics of this intricate recognition phenomenon. To this end, we have computationally simulated the dynamics and free energetics of domain II loop closing processes and identified a set of key amino acid residues of *Pf*RON2 helix which are essential for binding. The subsequent evaluation of the binding affinity of Ala-substituted *Pf*RON2 peptide ligands by surface plasmon resonance (SPR) and isothermal titration calorimetry (ITC) validates the relative importance of the residues in context. Together, the combination of computational and experimental investigation reveals that the domain II loop of *Pf*AMA1 is in fact responsible for arresting the *Pf*RON2 molecule from egress, K2027 and D2028 of *Pf*RON2 being the determinant residues for the capturing event. Our study provides a comprehensive understanding of the molecular recognition event between *Pf*AMA1 and *Pf*RON2, specifically in the post binding stage, which could potentially open up new avenues to drug discovery against malaria.

**TOC GRAPHIC:** 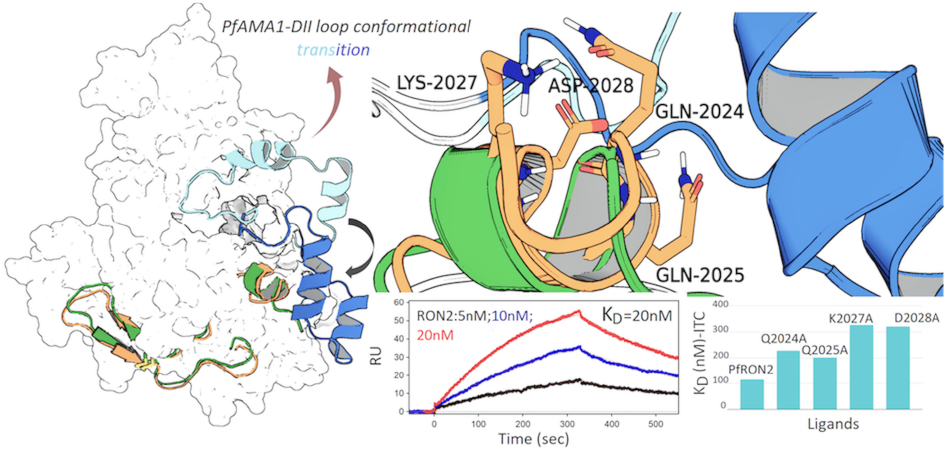

## INTRODUCTION

Malaria remains the dreaded disease with high mortality rates, mostly affecting populace of the economically challenged countries of the world. Despite the availability of approved therapeutic intervention strategies mortality caused by this disease poses a global health and economic concern, mostly due to continuous emergence of drug resistant strains (1, 2). One of the primal considerations for anti-malarial drug discovery is a thorough understanding of the malaria parasite invasion biology.

It is known that members of the phylum Apicomplexa, which include the malaria parasite *Plasmodium sp.*, exhibit common features in their invasion strategies despite their diverse host cell specificities and life cycle characteristics. Mechanistically, as pointed out by various research groups earlier, the formation of a moving junction between the membranes of the invading apicomplexan parasites and host cells is an essential step during the invasion process (3, 4). The moving junction in *Plasmodium sp.* comprises of two key parasite proteins; the surface protein apical membrane antigen 1 (AMA1) and its receptor, the rhoptry neck protein 2 (RON2). Reported crystal structures of the AMA1 have revealed that a partially mobile loop, termed domain-II (DII) loop, occupies part of a deep groove of the domain-I in the absence of its receptor RON2, apparently safeguarding the binding groove from any other possible polygamous recognition event (5, 6). During the moving junction formation, the DII loop gets replaced by the extracellular domain of RON2 (*Pf*RON_2021-2059_) letting the DII loop dangle freely as the RON2 interacts tightly within the binding pocket predominantly defined by an evolutionarily conserved hydrophobic trough. Atomistic details of *Pf*AMA1-*Pf*RON2_2021-2059_ recognition event, particularly, the role of the DII loop post RON2 binding, energetics related to it at large and the role of individual amino acid residues in specific, remain rather elusive till date.

Boulanger and co-workers, in one of their papers (4), provided significant clues pertaining to *Pf*AMA1-*Pf*RON2 interactions upon presence versus absence of the DII loop. They proposed that the DII loop contributes towards the kinetic locking of the *Pf*AMA1-*Pf*RON2_2021-2059_ complex. The 2-fold larger affinity and the 18-fold increased half-life of the *Pf*AMA1-*Pf*RON2_2021-2059_ complex in presence of the DII loop compared to the complex with truncated DII loop was attributed to the observed slower dissociation kinetics due to the restrained release of the *Pf*RON2 peptide in the former case (4). This report prompted us to investigate the atomistic details pertinent to the arrest of *Pf*RON2_2021-2059_ by the DII loop.

An overall understanding of this process remained difficult to address, as all AMA1-RON2 complex crystal structures available in the Protein Data Bank show missing electron densities (Lys351-Ala387) reiterating the intrinsic disorderness of the DII loop (7). Although a loop-closed crystal structure has been solved recently [PDB:6N87, (8)], it is still far from clear how the DII loop interacts with specific parts of the *Pf*RON2_2021-2059_ peptide or how the interim dynamics of the loop contributes to the overall stability of the complex. In the present work, we have addressed the questions mentioned above using atomistic molecular dynamics (MD) simulations (both unbiased and umbrella sampled, summing up to approximately 1.8 microsecond) coupled with biophysical experiments with systematic alanine substitution on the terminal helix of the *Pf*RON2_2021-2059_ peptide. We have used surface plasmon resonance (SPR) and isothermal titration calorimetry (ITC) to determine the binding affinity of chemically synthesized Ala-substituted *Pf*RON2_2021-2059_ analogues with AMA1 which helped us to experimentally verify the roles of key amino acid residues, originally identified by molecular simulations, that drive this recognition event.

## RESULTS

### Loop dynamics

Multiple repeats of atomistic simulations revealed that the DII loop reaches out towards the *Pf*RON2 peptide ligand (Fig.1A) in such a way that specific residues of the terminal helix of the *Pf*RON2 ligand interact with the loop as well as other adjacent residues of the AMA1 binding pocket. However, we focused only on the DII-loop-RON2 helix interactions to comprehend and quantify contributions from individual amino acids of the *Pf*RON2 helix which play dominant roles in this inter-molecular recognition event. We have computed distance profile between Ser373 of the DII loop and Gln2024 of the RON2 helix coupled with root mean square fluctuation (RMSF) to quantify the extent of conformational fluctuations (Fig. 1B). Distance profiles (Ser373-Gln2024) for all the trajectories have been given in Fig: S1 of supporting information. Residue level flexibility is one of the key metrics in understanding overall internal movement across simulation trajectories and is understood in terms of the RMSF. The observed RMSF of the whole molecule from all the simulation trajectories showed a highly stable nature, except the DII loop regions (residues 350 to 400) that showed the highest fluctuation compared to the rest of the residues. The flexible terminal residues were expected to fluctuate more than internal residues as they are destabilized in their local environment. The overall fluctuation ranged from 0.12 nm to 1.72 nm. As anticipated, the DII loop region was observed to fluctuate highest compared to other regions of the protein across all trajectories, depicting the movement of the DII loop (Fig: 1B and Fig: S3). Our study found evidences of intermittent association and dissociation of the RON2 helix and the DII loop, hinting at the role of DII loop in arresting the *Pf*RON2_2021-2059_ binding to *Pf*AMA1.

**Figure 1.**
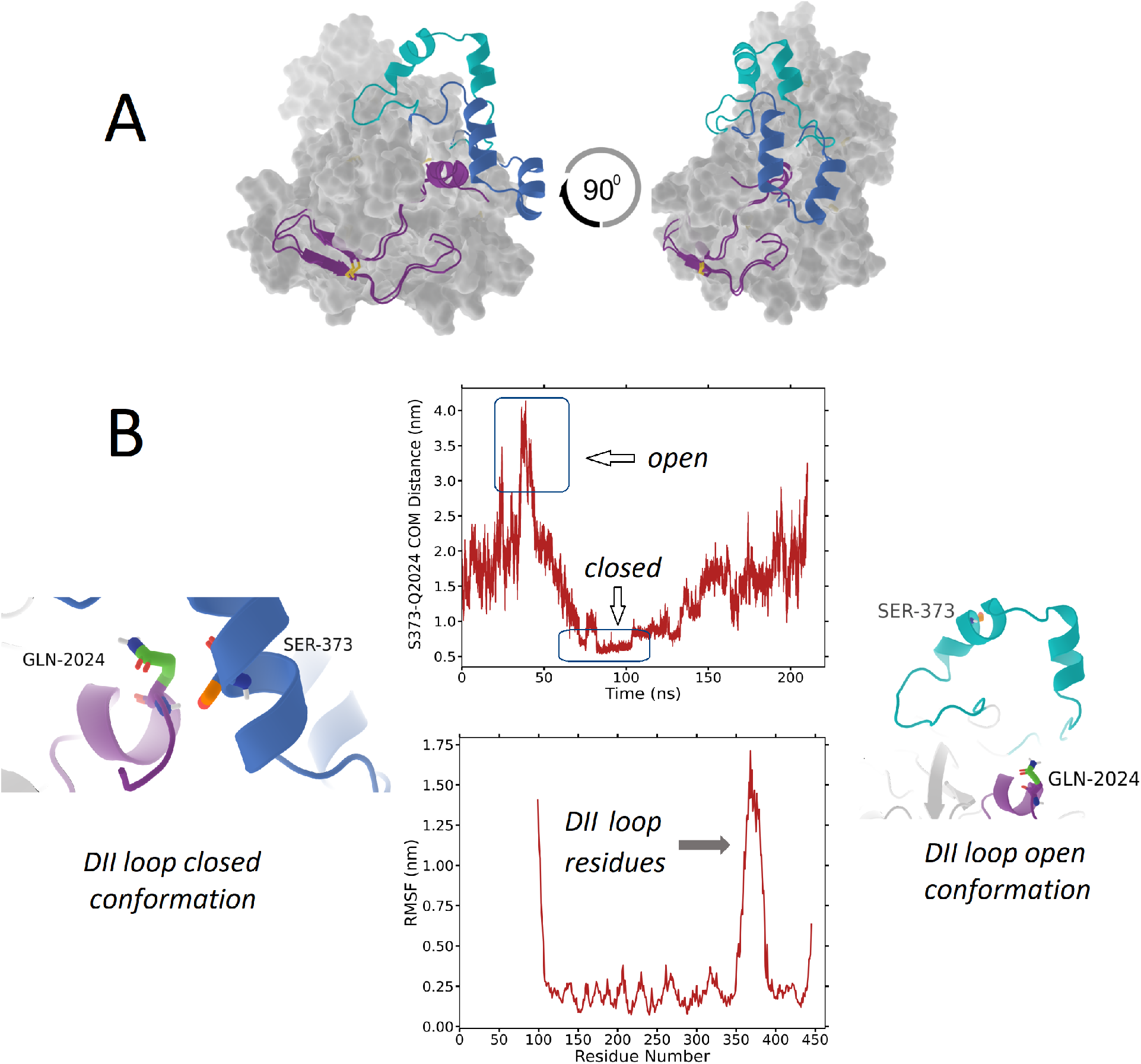
Loop closing event captured by molecular dynamics simulations. A. AMA1 is shown in surface representation; RON2 in purple; DII loop in cyan (open conformation) and deep blue (closed conformation). B. Centre of mass (COM) distance profile of Ser373 and Gln2024 of a representative trajectory (top) in closed (left) and open (right) conformation. The root mean square fluctuation (RMSF) plot of AMA1.

In the closed conformation we found that the Ile2022 formed hydrophobic contacts, intermittently, with the surrounding residues many of which are located at the DII loop. Most important residues making hydrophobic contacts with Ile2022 are Met374, Ile375, Phe385 and Ala378 of *Pf*AMA1 (Fig. 2).

**Figure 2.**
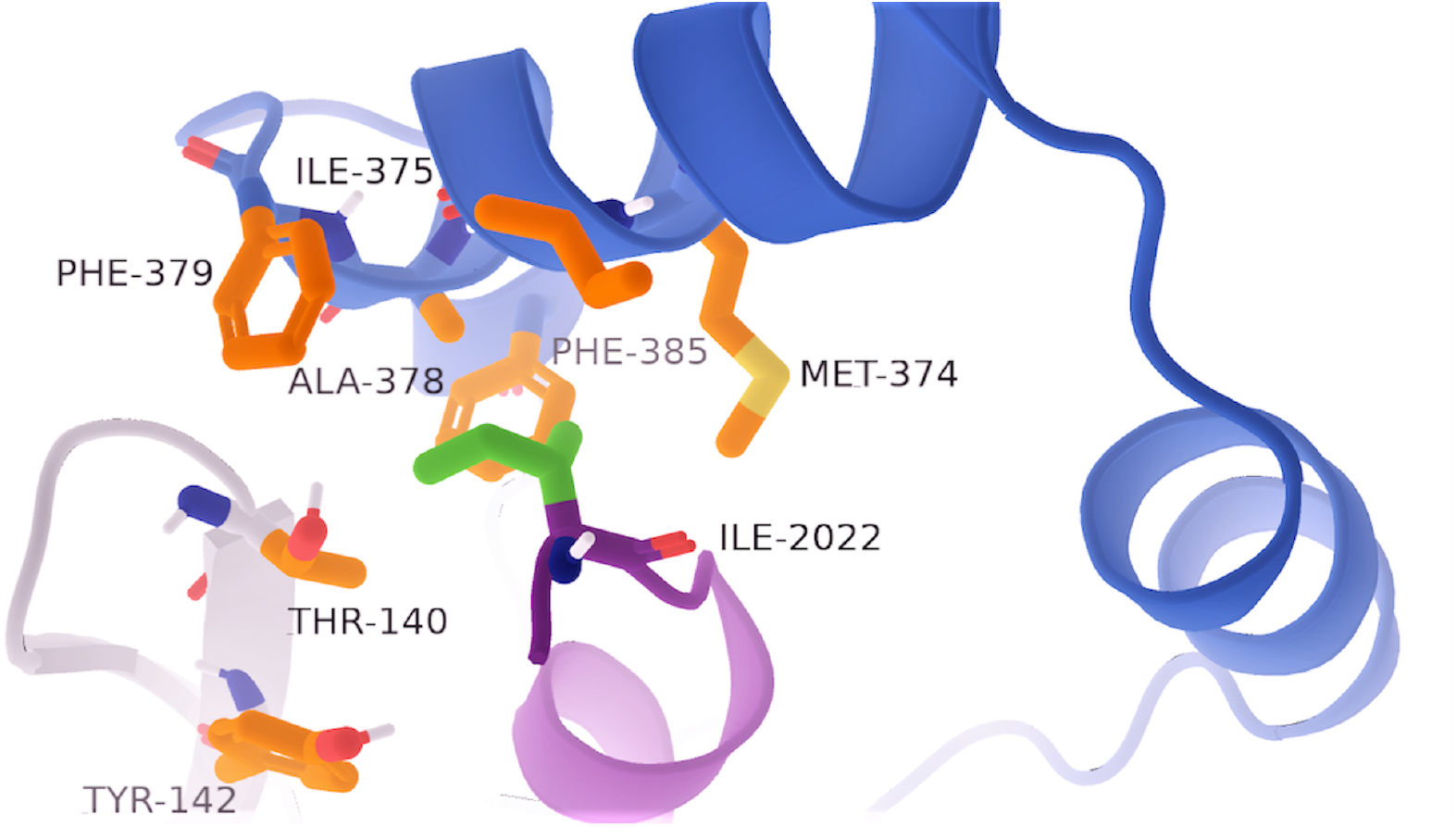
Hydrophobic pocket formed between Ile2022 and specific residues of AMA1 (RON2: purple, DII loop closed conformation: deep blue)

Most important interactions are, however, hydrogen bonds involving Gln2024, Gln2025, Lys2027 and Asp2028 which interact with various residues from the DII loop and adjacent areas of AMA1. Representative interactions between Lys2027 and DII loop residues are provided in Fig. 3 (detailed interactions are shown in Fig: S2).

**Figure 3.**
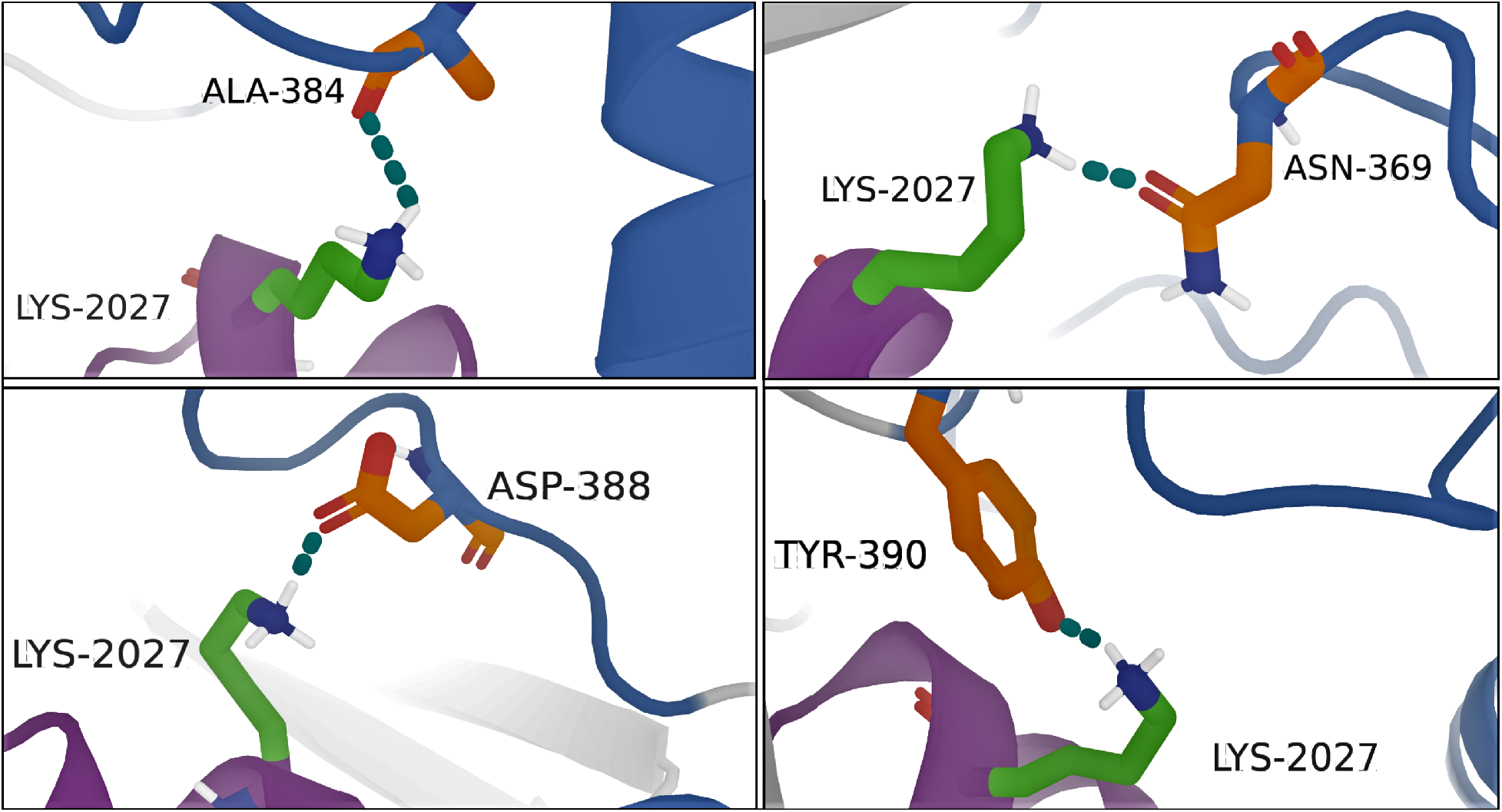
Hydrogen bonds formed between Lys2027 with representative residues of DII loop (detailed interactions are provided in SI-fig-2) (RON2: purple, DII loop closed conformation: deep blue, hydrogen bonds: teal)

The breathing motion of the DII loop and its interactions with the RON2 peptide observed in our MD simulations prompted us to examine the entire binding event in more quantitative manner. Towards this end, we envisioned that a meaningful free energetic insight on the DII loop closing event and its associated interactions with the alpha helix of RON2_2021-2059_ peptide can be achieved by simulating a free energy profile along an appropriate reaction coordinate. For this purpose, we used centre of mass distance between Ser373 and Gln2024 as the reaction coordinate. The free energy curve (see method section for details) indicated the formation of a favourable free energy basin when the residues mentioned in the collective variable came relatively closer (1.5Å) to each other and led to a state which we have defined as the closed conformation (Fig. 4). These results unambiguously stated that the DII loop possess high propensity of conforming to a state where it closes down towards AMA1 binding pocket and eventually arrests RON2 peptide from escaping.

**Figure 4.**
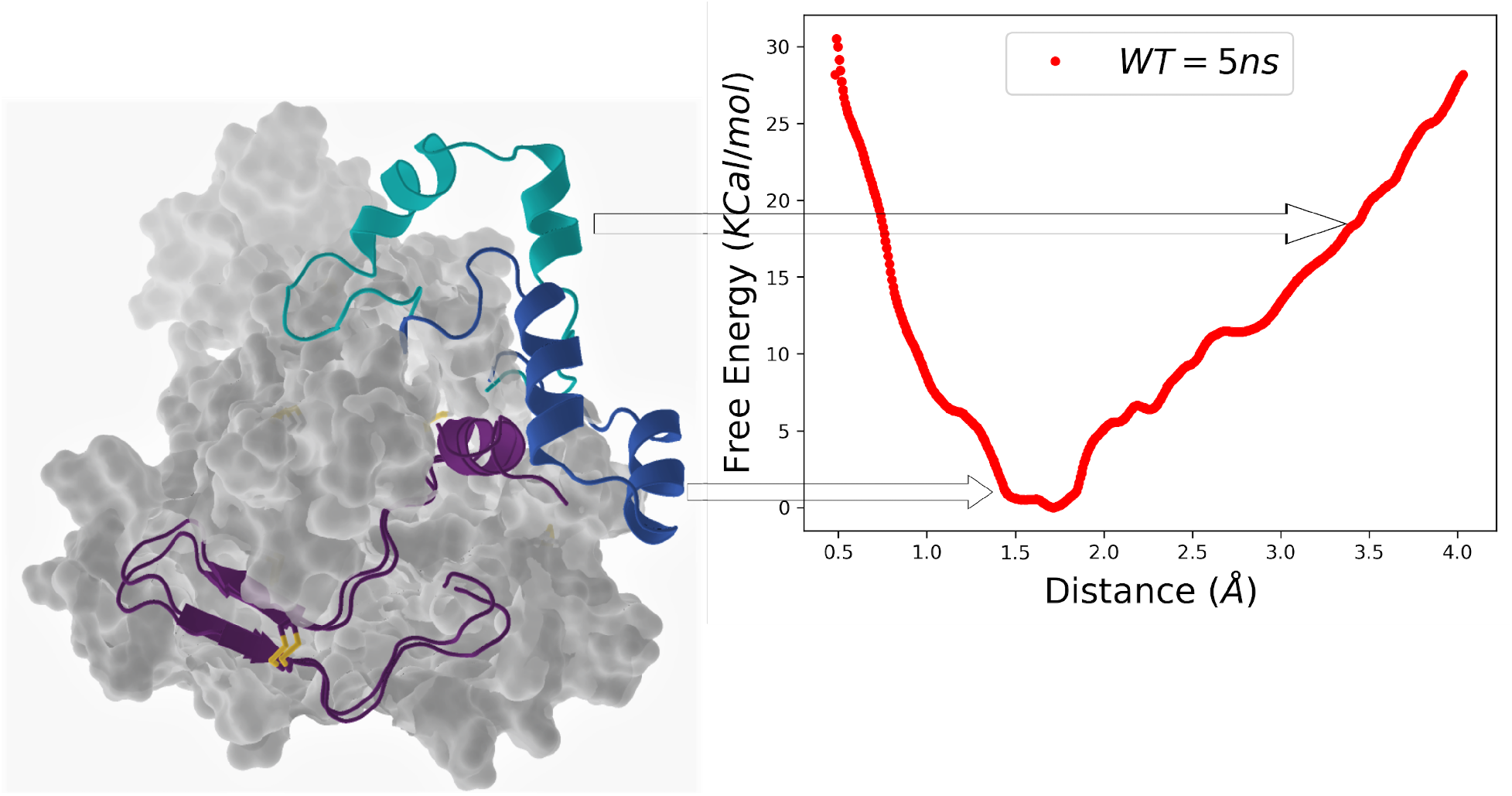
Free energetics of DII loop closing event. AMA1 represented in surface; RON2 in purple, DII loop in cyan (open conformation) and deep blue (closed conformation)

To further assess the status of the free energies of loop-closing upon mutation of specific amino acids, we performed independent free-energy calculations of a set of systems in each of which a single residue had been mutated to alanine. By doing a scan along a wide range of values of the chosen reaction coordinate (centre of mass distance between each of the residues Gln2024Ala, Gln2025Ala, Lys2027Ala and Asp2028Ala individually with Ser373), we obtained a measure of free energy of loop closing upon these mutations. A close inspection of these free energy curves revealed that the free-energy minima in all four mutants would appear at a relatively larger distance than that of wild-type, indicating that the mutations would result in a relatively open conformation compared to wild type system (Fig. 5). A possible reason of relatively open conformation in these mutants than the wild-type is presumably due to the lack of crucial hydrogen bonds in the former (Fig. 3). Additionally, we have observed strikingly different free energy profiles, especially for K2027A and D2028A, in which the open conformations are more stabilized than the loop-closed conformation. Our visual inspection of the simulation trajectories suggests that these two mutations make an unfavorable environment for the DII loop to close, due to the absence of suitable contacts with the alpha helix of RON2 peptide. As revealed in the aforementioned paragraphs, further experiments with ITC and SPR in the later sections also reiterate the relative importance of these residues.

**Figure 5.**
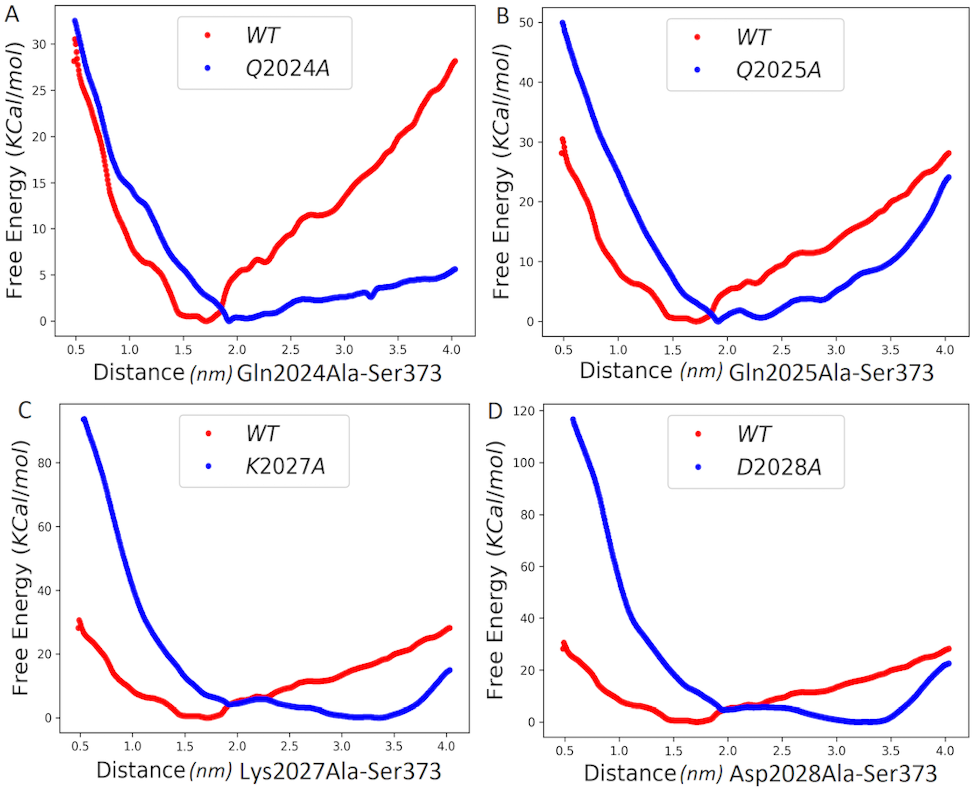
Free energy surfaces of Ala-substituted *Pf*RON2 analogues (A) Q2024A, (B) Q2025A, (C) K2027A and (D) D2028A compared with the wild type *Pf*RON2 (wild type: red and mutated: blue).

### *Pf*AMA1 expression from E. coli and in vitro refolding

Although *Pf*AMA1 contains three distinct domains, studies have revealed that domain I and II are responsible for the binding with the extracellular domain of *Pf*RON2(9). In order to understand the interactions of the *Pf*RON2 with domain II loop, we set out to express domain I and domain II of *Pf*AMA1 protein in Escherichia coli. We expressed the protein {*Pf*AMA1 (DI+DII)} in BL21 (DE3) RIL cells with an N-terminal His6-tag; so that the protein can be purified by immobilized metal affinity chromatography after extracting from the bacterial cell. The protein got expressed in the inclusion body of E. coli. After extraction from inclusion bodies, the reduced polypeptide was purified using Ni-affinity column followed by reverse phase HPLC and lyophilized. The lyophilized polypeptide was then refolded in vitro as described earlier (10). The final refolded protein was purified by size exclusion column chromatography to get rid of the other soluble misfolded proteins formed during the course of refolding. The freshly purified folded protein was used for SPR and ITC experiments to compare the binding affinity of the *Pf*RON2_2021-2059_ peptides and its Ala-substituted analogues.

### Chemical synthesis of the *Pf*RON2_2021-2059_ peptide and its analogues

To check the contributions of some of the key amino acid residues located in the *Pf*RON2 helix region in binding with the *Pf*AMA1 loop, we chemically synthesized the parent *Pf*RON2_2021-2059_ peptide as well as several other peptides where some of the key residues, which were computationally predicted to be responsible for binding with the DII loop region, were replaced by an alanine residue, one at a time. All peptides were synthesized by step-wise Fmoc SPPS using an automated peptide synthesizer. Lyophilized crude peptides were allowed to fold under air oxidation condition in 2M Gu.HCl, 100 mM Tris, pH 8 for 15h at room temperature to form one disulfide bond. Folded peptides were finally purified using reverse phase HPLC and lyophilized (Fig. 6)(10).

**Figure 6.**
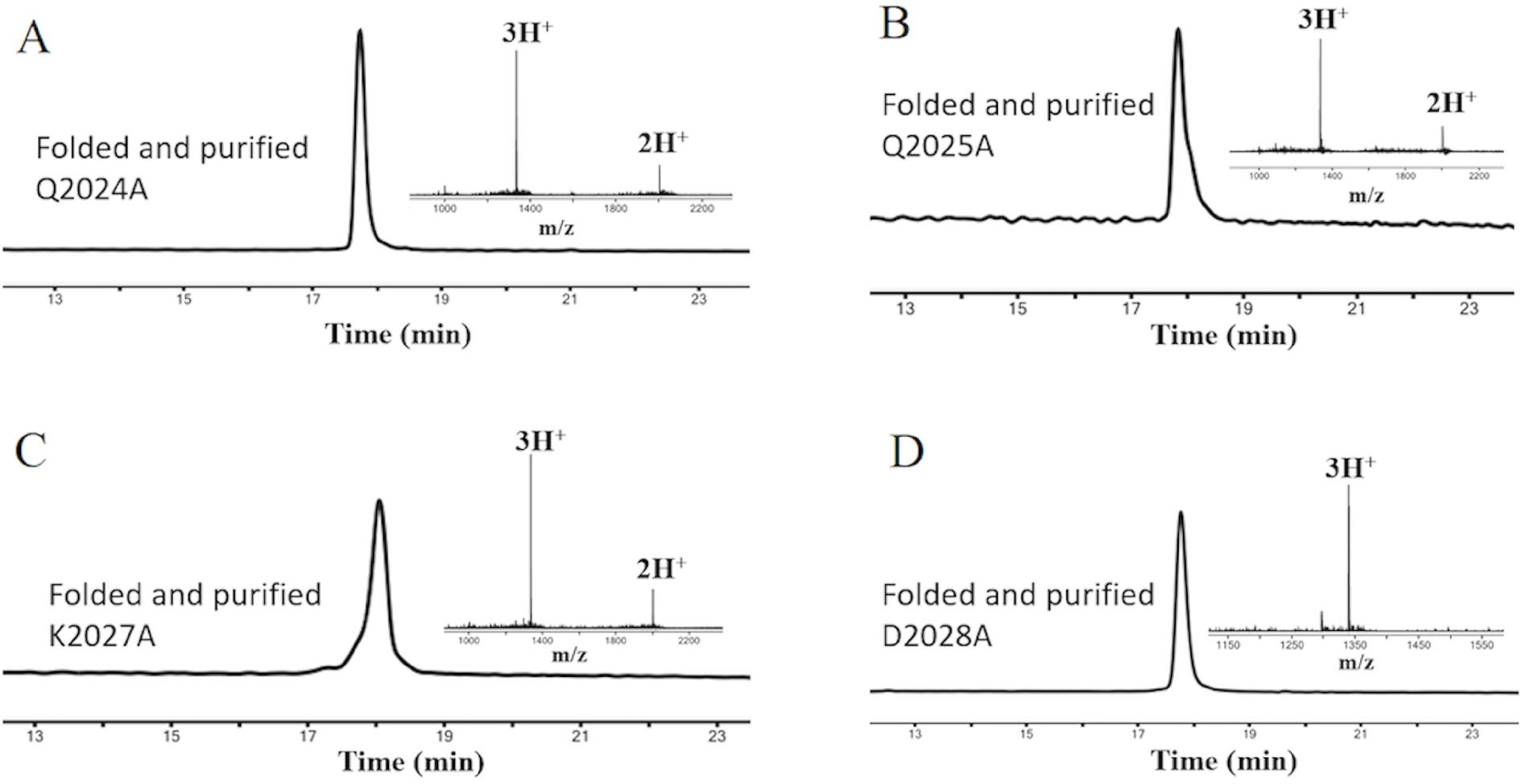
LC-MS chromatogram of the folded and purified *Pf*RON2 analogues. (A) Q2024A (calculated mass: 4003.97 Da, observed mass (most abundant isotopologue): 4003.99 ± 0.02 Da); (B) Q2025A (calculated mass: 4003.97 Da, observed mass (most abundant isotopologue): 4004.00 ± 0.02 Da); (C) K2027A (calculated mass: 4003.93 Da, observed mass (most abundant isotopologue): 4003.94 ± 0.01 Da); (D) D2028A (calculated mass: 4017.00 Da, observed mass (most abundant isotopologue): 4017.01 ± 0.02 Da).

### Binding study using SPR and ITC

Our goal was to evaluate the binding affinity of the Ala-substituted analogues of *Pf*RON2_2021-2059_ in order to identify their relative importance in binding with the DII loop. Therefore, we compared the binding profiles of the four peptide analogues (Q2024A, Q2025A, K2027A and D2028A) with *Pf*AMA1 with respect to *Pf*RON2_2021-2059_ peptide by SPR and ITC.

In SPR, we observed decrease in the association rate of RON2 analogues towards AMA1, while at the same time increase in the dissociation rate of the ligands from the AMA1-RON2 complex (Table 1). As a result, we observed a significant leap in the KD value of the Ala-substituted analogues when compared with the native RON2 ligand (KD= 22 nM)(10), indicating the weaker binding with AMA1 upon Ala substitution in the RON2 helix. We observed a consistent trend in the KD value between the substituted amino acid residues in SPR. We observed a significantly higher KD value of 899 nM and 228 nM for K2027A and D2028A analogues, respectively, indicating they are the two most important residues for the binding event; whereas, a KD of 141 nM and 75 nM were observed for the Q2024A and Q2025A analogues, respectively (KD values reported here are the average of three repetitions, Fig. 7 and Table-1). A similar trend in ΔG values was also evident from ITC indicating that the complex formed by the substituted RON2 peptide analogues with AMA1 is lesser stable than that of the AMA1-RON2 (Fig. 8, Table-1). The KD values of the analogues obtained from ITC also showed similar trend as SPR, which reveals that K2027 and D2028 are the most important residues in the binding phenomena followed by Q2024 and Q2025. The observed KD values from ITC for Q2024A, Q2025A, K2027A and D2028A, were 228 nM, 202 nM, 327 nM and 321 nM, respectively, whereas KD of native RON2 peptide was 115 nM as shown in fig-8 and Table-1. We note here that the data reported for the ITC are the average of two repeats due to the limitations of adequate quantity of the protein sample; whereas, for SPR it was the average of three repeat experiments. Moreover, as mentioned in Table 1, the error bars for the results obtained from ITC experiments are significantly higher, which is the limitation of the ITC technique. Hence, the data obtained from ITC should be interpreted carefully.

**Figure 7.**
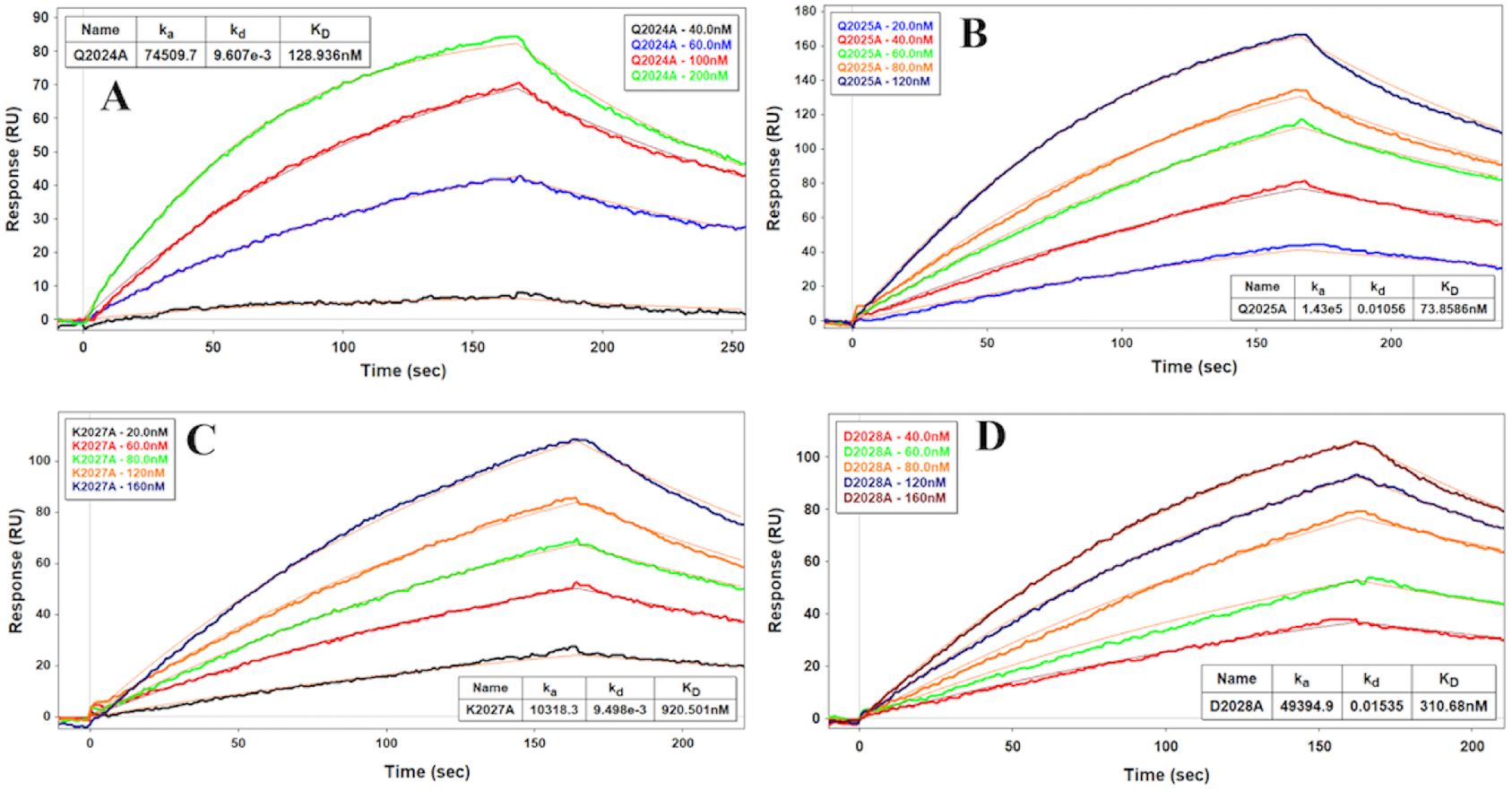
Representative SPR sensorgrams (one of the three repeats) of all alanine substituted peptides with *Pf*AMA1(DI+DII). (A) Q2024A (B) Q2025A (C) K2027A (D) D2028A.

**Figure 8.**
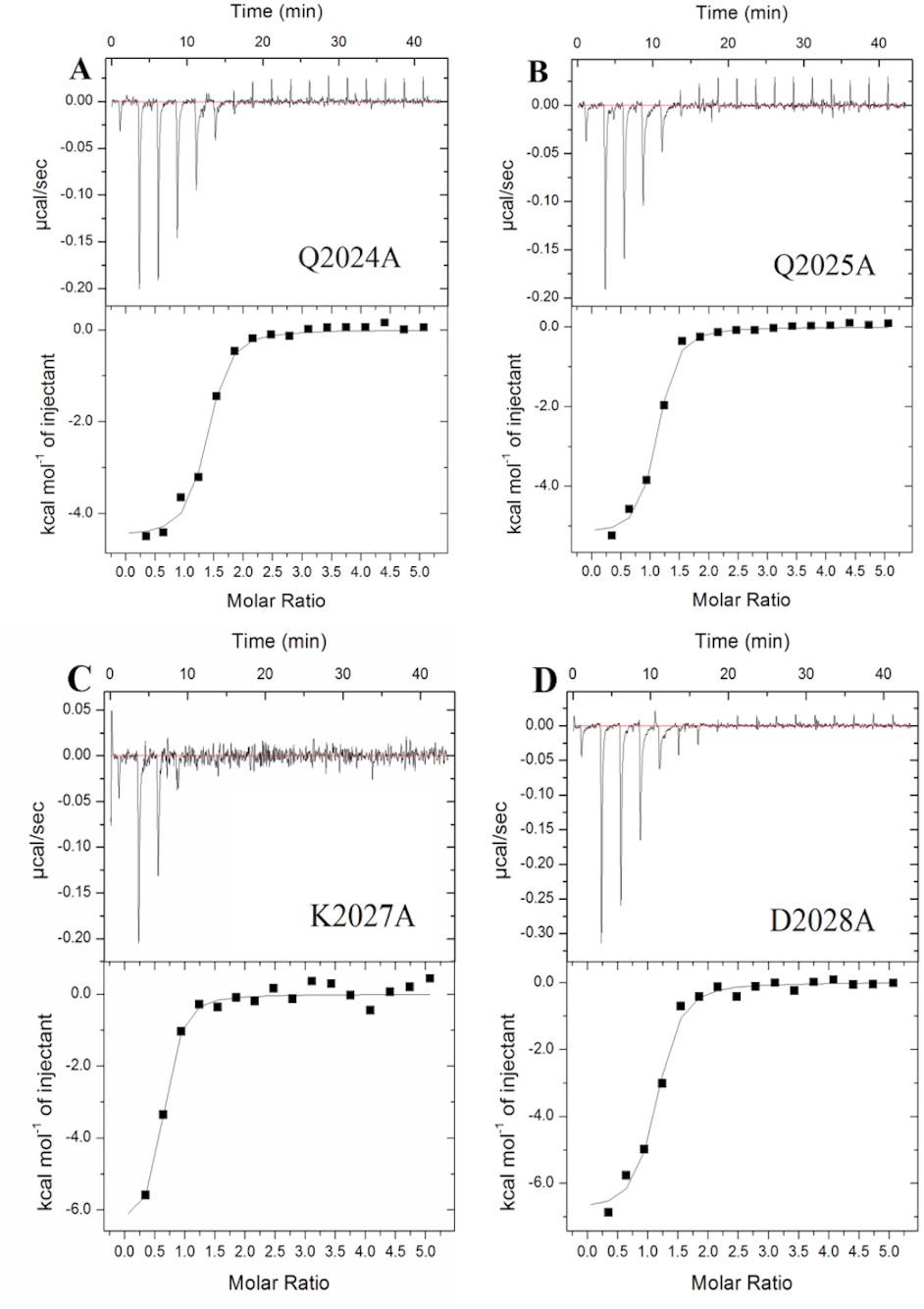
Binding profile of *Pf*AMA1 with all alanine substituted peptides in ITC experiment (representative one of the two repetitions). (A) Q2024A (B) Q2025A (C) K2027A (D) D2028A.

**Table 1.**
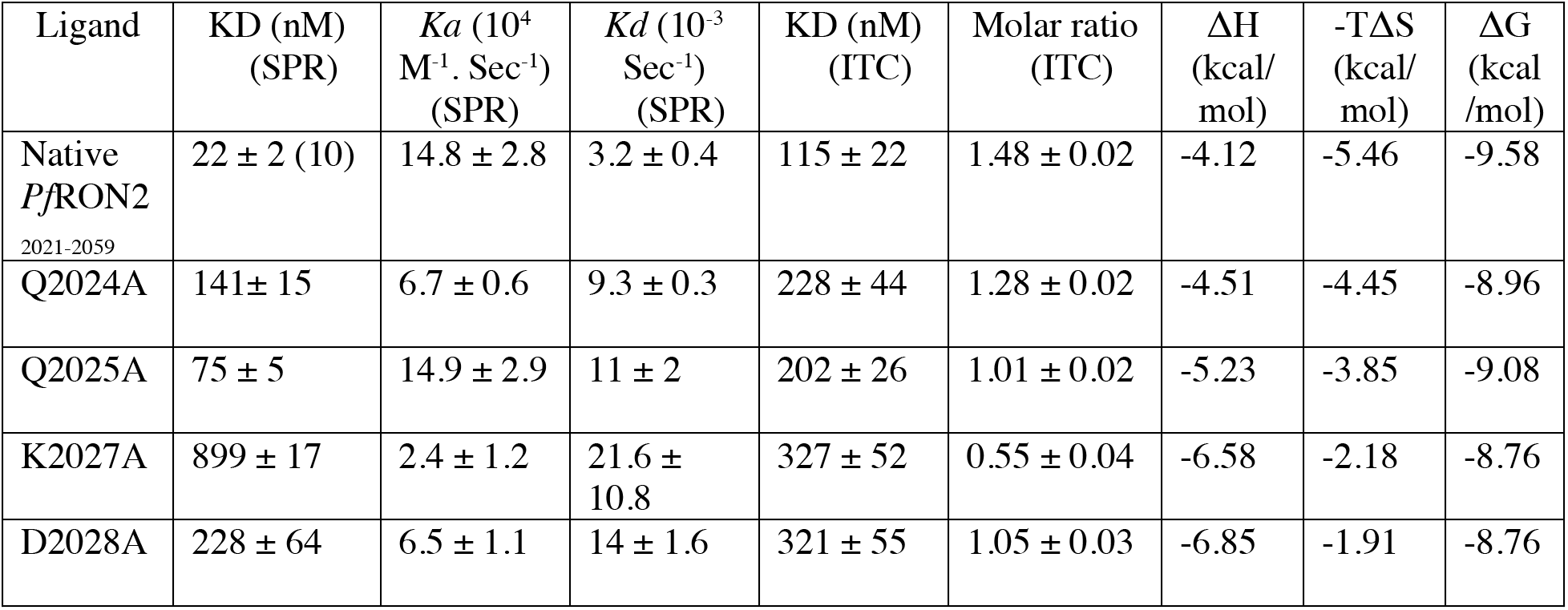
SPR (average of three repetitions; error represents the standard deviation of three experiments) and ITC (average of two repetitions) data comparing the binding affinity and thermodynamic parameters (one of the two repetitions; error represents the fitting error of the particular experiment) of the alanine substituted peptides with respect to the native *Pf*RON2_2021-2059_.

## DISCUSSION

Multiple repeats of atomistic simulations led us to infer that the DII loop, in effect, reaches out to the ectodomain of *Pf*RON2 in such a manner that the specific residues of the terminal helix of the *Pf*RON2_2021-2059_ interacts strongly with both DII loop and adjacent residues leading to the macromolecular arrest, which stops the egress of the bound *Pf*RON2 from the *Pf*AMA1 pocket. We have been able to decipher residue pairs which are responsible for the formation of free energetically favorable basins as a consequence of the closing of the DII loop after *Pf*AMA1-*Pf*RON2 binding event. As observed during MD simulations, Ile2022 makes hydrophobic contacts, intermittently, with Met374, Ile375, Phe385 and Ala378 leading to the formation of a cryptic hydrophobic pocket. However, the most important interactions are originated from Gln2024, Gln2025, Lys2027 and Asp2028 of the *Pf*RON2_2021-2059_ peptide which are involved in hydrogen bonding interactions with various residues from the DII loop and adjacent areas of the AMA1 (Fig. 3). Converged free energy calculations eventually led us to reach to a meaningful atomistic insight into the mechanism of DII loop closing and its interactions with the alpha helix of the *Pf*RON2 peptide. The free energy surface plot indicated formation of a favorable free energy basin when the residues mentioned in the collective variable came relatively closer, a state which we have defined as the closed conformation (Fig. 4).

To verify the importance of selected few most interactive residues, predicted from our computation data, we systematically substituted them with alanine. We observed that the loop closing was free energetically unfavorable with the chemically synthesized Ala-analogues, as the free energy dividend for bringing the loop closer from far off was more positive in Ala-substituted peptides (Fig. 5). According to our calculations, Lys2027 and Asp2028 were the most important residues from *Pf*RON2_2021-2059_ peptide contributing most in this intra-molecular recognition event. This claim was justified by both free energy simulations as well as the experimental data obtained from binding studies of the *Pf*RON2_2021-2059_ and its analogue peptides with *Pf*AMA1 by ITC and SPR experiments. In ITC, the average KD was observed to shift from 115 nM (*Pf*RON2_2021-2059_) to 327 nM and 321 nM in case of K2027A and D2028A, respectively (Table-1). However, in SPR experiment this KD shift was observed from 22 nM (*Pf*RON2_2021-2059_) to 899 nM and 228 nM for K2027A and D2028A, respectively (Table 1).

These results led us to believe that Lys2027 and Asp2028 are the most important residues that interact with the DII loop during the recognition event. This fact is further illustrated in Fig. 5 (C and D) where the free energy minima is found to be appearing far away compared to native *Pf*RON2_2021-2059_ (Fig. 4), which clearly explained the reluctance to let the event of loop closing happen upon substitutions at these positions. Similar trends were also noticed in case of other substitutions such as Q2024A and Q2025A where KD obtained from ITC was found to shift to 228 nM and 202 nM, respectively, as compared to 115 nM for *Pf*RON2_2021-2059_ and also the KD obtained from SPR was found to shift to 141 nM and 75 nM respectively as compared to *Pf*RON2_2021-2059_ which is 22 nM (Table-1). The combined results from molecular dynamics and systematic alanine substitution experiments further confirm that hydrogen bonds are accountable for the loop closing event to a larger extent than the hydrophobic transience exhibited by Ile2022 along with feeble hydrogen bonds contributed from its main chain.

Recent literature report (4) using ITC studies states that the free energy of binding of *Pf*RON2_2021-2059_ (ΔG=−10.6 kcal/mol) with *Pf*AMA1 in presence of DII loop is more favourable than with the AMA1 in absence of DII loop (ΔG = −9.6 kcal/mol), whilst witnessing a significant leap in KD (92.6±6.4 nM in the case of truncated DII loop versus 42.9 ±11.4 nM where the DII loop is intact). Although the DII loop thermodynamically stabilizes the binding of *Pf*RON2_2021-2059_ by a 2-fold larger affinity, it was observed by the authors to possess a particularly influential role on the kinetics by restraining the release of the PfRON2 peptide and increasing the complex half-life by 18-fold with a value of 900s (15 min) at 25°C for *Pf*AMA1-*Pf*RON2_2021-2059_ complex compared to 48s for ΔDII loop truncated *Pf*AMA1-*Pf*RON2_2021-2059_ complex (4). Contextually, a truncated *Pf*RON2 peptide (KDIGAGPVASCFTTRMSPPQQICLNSVVN), which also targets AMA1 but lacks a terminal helix, blocked invasion by preventing interactions of *Pf*AMA1 with the *Pf*RON2 (KD=520±74 nM; PDB-3SRI). This fact also supports our understanding, with caveats though, as the free energy of binding of the truncated *Pf*RON2 peptide has not been reported so far (7). In another recent work, pertaining to the design of β-hairpin peptides (Cys2037-Cys2049 patch), which completely excludes the alpha helix motif also showed diminished functional activity (KD=57±12 *μ*M, (11)). Therefore, it was evident that both DII loop and the alpha helix of RON2 are absolutely necessary to exhibit an effective binding event. In the current study, while investigating the quantitative role of individual amino acid residue of RON2 helix in the binding event by systematically altering the helix residues with alanine keeping the beta-loop intact, computational results revealed that the primary mode of contact of RON2 with AMA1 is significantly unperturbed. However, substitutions at the particularly designated positions, as hinted from multiple repeats of molecular simulation studies, led to the disruption of interactions of the RON2 helix with AMA1. In case of these Ala-analogues of *Pf*RON2_2021-2059_, the structural perturbations caused by Ala substitutions led to a faster egress of RON2 from the AMA1 binding pocket. This fact was further bolstered from the multiple replicas of binding assay measured by SPR, where we have observed quicker dissociation rates in case of the alanine substituted *Pf*RON2_2021-2059_ analogues compared to the native one. It was observed that K2027A (*kd* = 21.6 × 10^−3^ Sec^−1^) and D2028A (*kd* = 14.0 × 10^−3^ Sec^−1^) dissociate faster followed by Q2024A (*kd* = 9.3 × 10^−3^ Sec^−1^) and Q2025A (*kd* = 11.0 × 10^−3^ Sec^−1^); whereas, in case of native *Pf*RON2_2021-2059_ the dissociation rate was least (*kd* = 3.2 × 10^−3^ Sec^−1^) among all five cases (Table 1). This observation indicates the suitability of wild type residues in those particular positions as a decisive metric towards arrest of *Pf*RON2_2021-2059_ to the AMA1 binding pocket. Moreover, repeated ITC experiments on the substituted *Pf*RON2_2021-2059_ peptides showed significant change in the free energy of binding against *Pf*AMA1(DI+DII), making the substitutions unfavourable for the AMA1-RON2 complex formation. The free energy of binding decreases the most in case of K2027A (ΔG = −8.76 kcal/mol) and D2028A (ΔG = −8.76 kcal/mol) compared to *Pf*RON2_2021-2059_ (ΔG = −9.58 kcal/mol) further indicating the importance of these two residues. We have observed a jump in the free energy of binding in case of the other two substitutions as well (ΔGQ2024A = −8.96 kcal/mol, ΔGQ2025A = −9.08 kcal/mol) with respect to the native *Pf*RON2_2021-2059_, although relative importance is lesser than that of K2027A and D2028A.

## MATERIALS AND METHODS

### MD simulations and the potential of mean force (PMF) calculations using umbrella sampling

We performed MD simulations using the GROMACS code (version 2019)(12, 13). For simulations we used the CHARMM36 force field (14) with TIP3P water model (15). Charmm-gui(16) web-server was used to prepare the systems including the construction of the loop. Coordinates from PDB:3ZWZ were used as a starting structure for simulations. A cubic box type with dimensions of 10.10 nm^3^ was used. Then, the box was solvated with 30331 TIP3P water molecules and the system was neutralized with potassium chloride counter ions in a concentration of 0.15 mol/lit. Total number of atoms in the system was 97154. The steepest descent algorithm (17) was applied to optimize the system to its lowest possible energy. A position restraint of 1000 kJ mol^−1^ nm^−2^ was applied to fix the positions of both ligand and protein atoms and the whole system was equilibrated under the NVT ensemble through 100 ps time during which the V-rescale thermostat was used to regulate the temperature close to 310 K. Following the NVT step, the system was equilibrated under the NPT ensemble through 100-ps time in such a way that the pressure of the system was stabilized to 1 atm. Multiple repeats of production MD runs were performed using 2 fs as timestep with usage of leap-frog integrator, differing in the assignment of initial velocity seeds on the well equilibrated system at the temperature of 310 K and the pressure of 1 atm. The simulations were initially continued to capture the entire breathing motion of the loop, i.e., from open conformation to closed conformation and again to open conformation and few of the simulations were continued till the event of loop closing were observed. In the later stages, all these trajectories were concatenated to calculate the cumulative properties like hydrogen bonds between DII loop residues and RON2 helix. Hydrogen bonds were calculated as criteria specified by Gromacs (https://manual.gromacs.org/documentation/current/reference-manual/analysis/hydrogen-bonds.html) where hydrogen bonds are determined based on cutoffs for the angle Hydrogen - Donor - Acceptor and the distance Donor-Acceptor. OH and NH groups are regarded as donors, O is an acceptor always and N is an acceptor by default. The Particle-mesh Ewald (PME) algorithm(18) was used to calculate the long-range electrostatic contributions and the length of all covalent bonds was constrained using the LINCS algorithm(19), a faster algorithm in comparison to the SHAKE algorithm (20). An analytical algorithm, called SETTLE, for resetting the positions and velocities to satisfy the holonomic constraints on the rigid water model was used (21). Upon termination of the simulation, the protein was backed and centered in the box along with removal of the periodic boundary condition from the trajectory.

The potential of mean force for dynamics of the DII loop movement with reference to RON2 peptide was constructed using umbrella sampling MD simulations. Umbrella sampling (22) is a very popular technique for PMF calculation to study binding-unbinding processes related to protein-ligand and protein-protein interactions (23–25). The binding free energy can also be extracted from the obtained PMF. The umbrella sampling is a non-Boltzmann sampling where an extra term is added to the hamiltonian for sampling to explore the collective variable (CV) space. In practice, CV space is divided into a series of windows and an external potential, usually harmonic, is applied to keep the distribution peaked within a region of the order parameter. As CV, we used centre of mass distance between two amino acids (Ser373 and Gln2024) as deduced from the unbiased simulations. 50 umbrella windows were created along this distance at 0.5 Å intervals with a force constant (k) of 500kcal/mol/Å^2^ until it indicated an overlapping histogram of positional distributions. The umbrella sampling calculations were done for 5 ns in the wild type system and 5 ns for the alanine substituted analogues for each window. Finally, we used weighted histogram analysis method (WHAM)(26, 27) which combines the individual PMFs optimally to weigh the individual distributions to obtain the free energy profile.

### Expression, purification and refolding of *Pf*AMA1(DI+DII)

The expression, purification and refolding of the *Pf*AMA1(DI+DII) was performed following the published procedure (10). In brief, the expression plasmid encoding the sequence of 3D7 *Pf*AMA1(DI+DII) (104-438) was obtained commercially from GenScript (NI, USA) having N-terminal hexa-His tag along with one TEV-protease cleavage site. The protein was expressed in Escherichia coli BL21 (DE3) RIL cells and purified through Ni-affinity column followed by reverse phase HPLC purification and lyophilization. Lyophilized protein was then refolded and purified by size exclusion chromatography as described earlier (10). Protein containing fractions were pooled and used for the binding assay by SPR and ITC.

### Synthesis and oxidative folding of RON2 and other alanine substituted peptides

The 39-mer extracellular domain of RON2 (Asp2021-Ser2059, *Pf*RON2_2021-2059_) was synthesized on 2-chlorotritylchloride resin using Fmoc protected amino acids by solid phase peptide synthesis (SPPS) at a single stretch using an automated peptide synthesizer (Tribute UV-IR, Protein Technologies Inc. USA). After cleaving the peptide from the solid resin support, the crude peptide was dissolved in 6M guanidine hydrochloride (Gu.HCl), centrifuged, filtered with 0.2 micron filter and slowly diluted with Tris buffer to get the ultimate folding condition of 2M Gu.HCl and 100 mM Tris (pH 8). To form the disulfide bond, the peptide was incubated under air oxidation condition for 15h. The folded peptide was purified by reverse phase HPLC and lyophilized. The other Ala substituted analogues Q2024A, Q2025A, K2027A and D2028A) were also synthesized adapting the same protocol described above followed by oxidative disulfide formation, purification by reverse phase HPLC and lyophilization. All the lyophilized peptides were reconstituted in running buffer (10 mM phosphate buffer saline, pH 7.4, 0.005% Tween-20, 40 μM EDTA) for SPR experiments and in elution buffer (20 mM Tris pH 7.8, 100 mM NaCl) of size exclusion chromatography for ITC experiments to study the binding against *Pf*AMA1(DI+DII).

### Binding affinity of peptides with AMA1 using surface plasmon resonance

BI-4500AP SPR instrument was used to measure the binding of the *Pf*RON2_2021-2059_ peptides and its alanine substituted analogues with *Pf*AMA1. 2.4 μM freshly refolded His-tagged AMA1 protein was immobilized at pH 7.8 over a Nickel-NTA chip while 10 mM phosphate buffer saline, pH 7.4 with 0.005% Tween-20 was used as the running buffer. Three flow cells were used for the immobilization of the AMA1 protein using ‘G-inject’ function to get gradually decreasing or increasing level of immobilization. The reference flow-cell was created by injecting only the running buffer. Before passing through the analytes over the chip, 40 μM EDTA was added in the running buffer to minimize non-specific interactions. A series of concentrations of a particular analyte dissolved in the running buffer (10 mM Phosphate buffer saline, pH 7.4 with 0.005% Tween-20 and 40 μM EDTA) were then passed through all the four flow-cells at a constant flow rate of 30 μl/min to obtain the association curve. After 360 sec (for *Pf*RON2_2021-2059_) and 150 sec (for Ala substituted peptides due to their weaker binding propensity), the injection of the peptide analytes was stopped and the chip was allowed to regenerate in the running buffer at the same flow rate to capture the dissociation event. The final sensorgram was generated by subtracting the reference flow cell from the ligand flow cell and the resulting data was fitted with 1:1 Langmuir adsorption binding isotherm using the BI-data analysis software. For all analytes, the binding experiments were performed in three repeats using the same batch of refolded *Pf*AMA1(DI+DII) protein and reported as an average of the three data.

### Binding affinity of peptides with AMA1 using isothermal titration calorimetry

MicroCaliTC200 instrument was used to check the binding affinity of the peptides with the *Pf*AMA1 protein. 200 *μ*l of the 10 *μ*M *Pf*AMA1 was loaded into the ITC sample cell and the protein was titrated with 2 *μ*l of the ligand (peptide) for total 17 injections. Ligands were dissolved in 20 mM Tris and 100 mM NaCl (pH 7.8), the same buffer in which the AMA1 protein was eluted. In optimized conditions, *Pf*RON2_2021-2059_ concentration was kept as 200 *μ*M while the other Ala substituted peptide concentrations were kept at 300 *μ*M due to their weaker binding affinity towards *Pf*AMA1 protein as compared to *Pf*RON2_2021-2059_. Heat of dilution data was recorded in Tris-NaCl buffer for each peptide and final data was obtained by subtracting this data from the raw data. The data fit was obtained using Origin software (Microcal) and dissociation constant was calculated for each ligand using one site model. The binding study for each peptide was repeated twice for reproducibility against freshly refolded *Pf*AMA1(DI+DII) and KD values were reported as average of the two.

## CONCLUSIONS

A fundamental understanding of the mode of interactions between the two important Plasmodium falciparum protein partners like AMA1-RON2 is absolutely necessary to design potential therapeutics against the dreaded disease like malaria. To that end, the current work undertook both computational as well as experimental binding studies with systematically substituted alanine analogues of *Pf*RON2_2021-2059_ in order to understand, in detail, how the *Pf*RON2_2021-2059_ interacts with AMA1, with special emphasis on the role of DII loop in the due process of the recognition event. From the detailed analysis of the computational as well as the experimental data, we have found a direct correlation between the DII loop and the alpha helix of the *Pf*RON2_2021-2059_ that influence each other leading to the formation of a stable complex. We have not only identified key binding residues of the alpha helix of the *Pf*RON2_2021-2059_ but also many other important molecular interactions responsible for carrying out this binding event. The current report also reiterates previous findings on specific kinetic aspects, like egress rates, which are found to be significantly perturbed by the Ala substitutions of specific residues on the N-terminal alpha helix of the *Pf*RON2_2021-2059_. In summary, the current work provides novel insights into key interactions between *Pf*AMA1 and *Pf*RON2 proteins which will further assist in designing potential RON2 based AMA1 inhibitors.

## Supporting information

Supporting Informations

## Supporting Information

A Supporting Information file containing Figure S1-S3 and Table S1-S3 has been included (file type PDF).

## Author Contributions

J.M. and K.M. conceptualization; S.S., A.B., J.M., and K.M. designing experiments; S.S., and A.B. data curation; S.S., A.B., J.M., and K.M. analyzing data; S.S., and A.B. writing – original draft; S.S., A.B., J.M., and K.M. writing – review and editing; J.M., and K.M. supervision and funding acquisition. All authors have given approval to the final version of the manuscript.

## ACKNOWLEDGMENT

This work was supported by the DBT/Wellcome Trust India Alliance (grant no. IA/I/15/1/501847) to K.M. and the Department of Atomic Energy, Government of India, under Project Identification No. RTI 4007 to J.M. and K.M. The authors acknowledge the biophysics core facility for the ITC experiments and computational core facility of TIFR Hyderabad. The authors thank Dr. Sreejith Rarankurusi for assisting with ITC experiments, Satyabrata Bandyopadhyay and Souvik Sadhukhan for their help with umbrella sampling and python plotting.

