## Supporting Informations for "Computational and experimental studies of the breathing motion of a protein loop: implications in *Pf*AMA1-*Pf*RON2 late-stage binding event"

[‡] Equal contribution

---

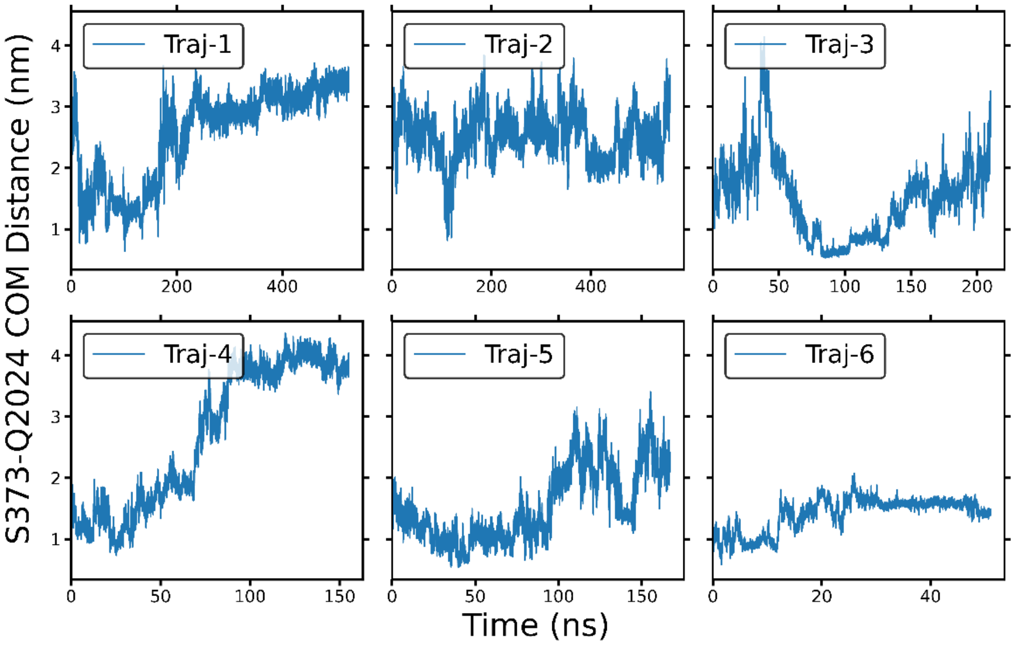


**Figure S1**: Distance profiles between Ser373 and Gln2024 in all trajectories

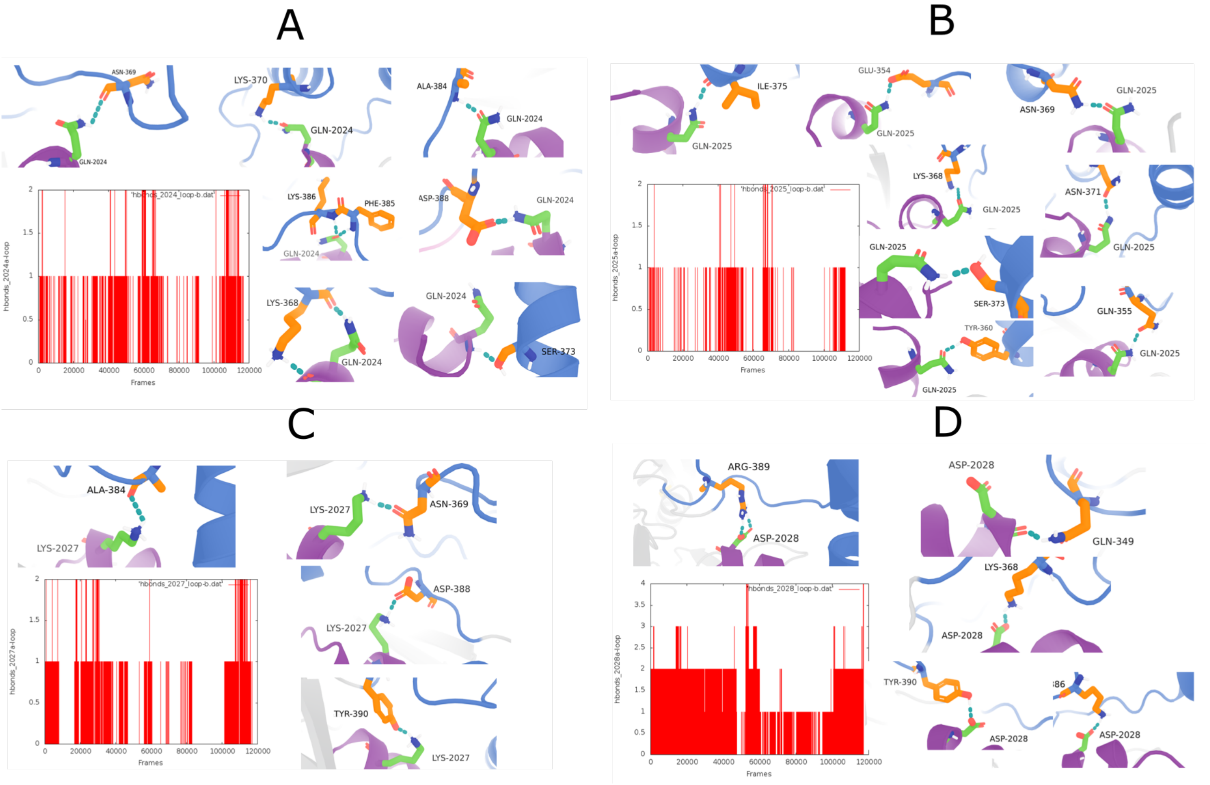


**Figure S2**. Detailed hydrogen bond interactions between Gln2024 (A), Gln2025 (B), Lys2027 (C), Asp2028 (D) with representative residues of DII loop.

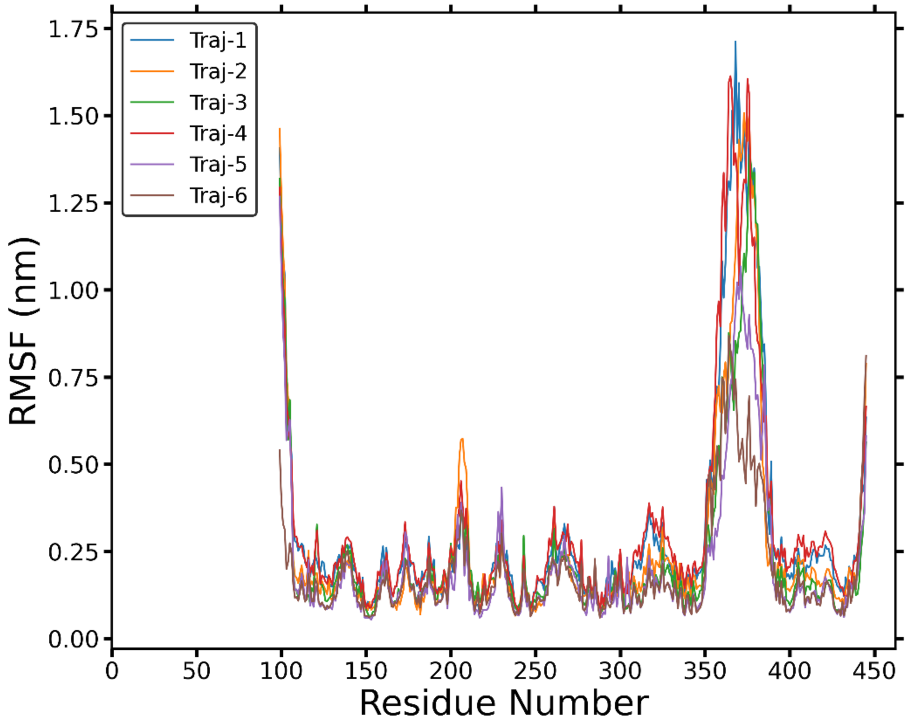


**Figure S3:** Superposition of the RMSF plot from all trajectories

**Table S1.** Trend in change of affinity upon alanine substitution experiments.

| Ligands | Fold increase in $K_D$ (SPR) | Fold increase in $K_D$ (ITC) |
|---|---|---|
| *Pf*RON2$_{2021\text{-}2059}$ | NA | NA |
| K2027A | 41.07 | 2.84 |
| D2028A | 10.40 | 2.78 |
| Q2024A | 6.44 | 1.98 |
| Q2025A | 3.42 | 1.75 |

**Table S2.** Summary of results obtained from multiple repeats of SPR experiments.

| Ligand | Repetitions | $K_a$ ($M^{-1}$. $Sec^{-1}$) | $K_d$ ($Sec^{-1}$) | $K_D$ (nM) |
|---|---|---|---|---|
| **Q2024A** | Set-1 | 74509.7 | 9.61E-03 | 129 |
| | Set-2 | 61882 | 9.26E-03 | 150 |
| | Set-3 | 60024.9 | 9.68E-03 | 161 |
| | Set-4 | 71030.5 | 8.82E-03 | 124 |
| **Q2025A** | Set-1 | 1.43E+05 | 0.01056 | 74 |
| | Set-2 | 1.58E+05 | 0.01065 | 67 |
| | Set-3 | 1.87E+05 | 0.01501 | 80 |
| | Set-4 | 1.06E+05 | 8.22E-03 | 77 |
| **K2027A** | Set-1 | 10318.3 | 9.50E-03 | 920 |
| | Set-2 | 22201.6 | 0.01951 | 879 |
| | Set-3 | 4.00E+04 | 0.03582 | 896 |
| **D2028A** | Set-1 | 49394.9 | 0.01535 | 311 |
| | Set-2 | 6.93E+04 | 0.01515 | 219 |
| | Set-3 | 7.77E+04 | 0.01193 | 154 |

**Table S3.** Summary of results obtained from two repeats of ITC experiments.

| | Set-1 | | | | | Set -2 | | | | |
|---|---|---|---|---|---|---|---|---|---|---|
| Ligand | $K_D$ (nM) | Molar ratio | ΔH (kcal/mol) | -TΔS (kcal/mol) | ΔG (kcal/mol) | $K_D$ (nM) | Molar ratio | ΔH (kcal/mol) | -TΔS (kcal/mol) | ΔG (kcal/mol) |
| Q2024A | 271.7 | 1.28 ± 0.02 | -4.51 | -4.45 | -8.96 | 184.2 | 1.69 ± 0.04 | -3.33 | -5.88 | -9.20 |
| Q2025A | 227.8 | 1.01 ± 0.02 | -5.23 | -3.85 | -9.08 | 175.4 | 0.99 ± 0.03 | -5.61 | -3.61 | -9.22 |
| K2027A | 378.8 | 0.55 ± 0.04 | -6.58 | -2.18 | -8.76 | 274.7 | 0.70 ± 0.01 | -6.00 | -2.96 | -8.95 |
| D2028A | 375.9 | 1.05 ± 0.03 | -6.85 | -1.91 | -8.76 | 265.3 | 0.85 ± 0.02 | -9.25 | 0.27 | -8.97 |
